# Alzheimer’s disease risk SNPs show no strong effect on miRNA expression in human lymphoblastoid cell lines

**DOI:** 10.1101/367318

**Authors:** Inken Wohlers, Colin Schulz, Fabian Kilpert, Lars Bertram

**Affiliations:** Lübeck Interdisciplinary Platform for Genome Analytics (LIGA), Institutes of Neurogenetics and Cardiogenetics, University of Lübeck, Lübeck, Germany; Centre for Lifespan Changes in Brain and Cognition, Dept of Psychology, University of Oslo, Oslo, Norway

**Author notes:** Correspondence to Prof. Lars Bertram, Lübeck Interdisciplinary Platform for Genome Analytics (LIGA), Institutes of Neurogenetics and Cardiogenetics, University of Lübeck, Maria-Goeppert-Str. 1, 23562 Lübeck, Germany.

**Keywords:** miRNA, gene expression, genetic association, GWAS, eQTLs, Alzheimer’s disease

## Abstract

The role of microRNAs (miRNAs) in the pathogenesis of Alzheimer’s disease (AD) is currently extensively investigated. In this study, we assessed the potential impact of AD genetic risk variants on miRNA expression by performing large-scale bioinformatic data integration. Our analysis was based on genetic variants from three AD genome-wide association studies (GWAS). Association with miRNA expression was tested by expression quantitative trait loci (eQTL) analysis using next-generation miRNA sequencing data generated in lymphoblastoid cell lines (LCL). While, overall, we did not identify a strong effect of AD GWAS variants on miRNA expression in this cell type we highlight two notable outliers, i.e. miR-29c-5p and miR-6840-5p. MiR-29c-5p was recently reported to be involved in the regulation of *BACE1* and *SORL1* expression. In conclusion, despite two exceptions our large-scale assessment provides only limited support for the hypothesis that AD GWAS variants act as miRNA eQTLs.

## 1 Introduction

Alzheimer’s disease (AD) is the most common age-related neurodegenerative disease characterized by progressive cognitive decline eventually leading to dementia. While it is established that genetic factors play a crucial role in determining liability to both monogenic as well as polygenic forms of AD, the underlying functional mechanisms – in particular those driving polygenic AD – hitherto remain only poorly understood. One such mechanism may involve the (dys-)function of microRNAs (miRNAs). MiRNAs are small, non-coding RNAs that post-transcriptionally alter messengerRNA (mRNA) expression and consequently protein abundance by their semi-complementary binding to specific mRNA targets. Like the expression of protein-coding transcripts (i.e. mRNAs) a substantial portion of miRNA expression is genetically controlled by alleles of both common and rare DNA sequence variants. Here, we systematically investigated whether and which of the currently established AD risk variants exhibit allele-specific effects on miRNA expression utilizing next-generation sequencing-based miRNA expression data in human lymphoblastoid cell lines previously generated by the GEUVADIS consortium. For two miRNAs whose expression was found to be at least partially regulated by AD associated variants, we performed additional analyses to characterize their potential role in AD pathogenesis.

## 2 Methods

Data integration is based on genome-wide AD association results retrieved from publicly available GWAS summary statistics from the International Genomics on Alzheimer’s Disease Project (IGAP (2013)). These data were supplemented by index SNPs from two subsequently reported GWAS (Jansen et al. (2018) and Kunkle et al. (2018)). MiRNA eQTL analysis was performed using small RNA sequencing data generated in 345 independent lymphoblastoid cell lines by the GEUVADIS consortium (Lappalainen et al. (2013)). GWAS and miRNA expression data were integrated using a variety of methods including a data integration pipeline (ADmiReQTL) developed by our group and two recently developed tools for assessing the relationship between genetic association and gene expression data (JLIM (Chun et al. (2017)) and TWAS/FUSION (Gusev et al. (2018))).

### GWAS data

For our genome-wide analyses we used the data released by the IGAP consortium which represents the largest currently available set of AD GWAS summary statistics (IGAP (2013)). For the analyses of this paper, we utilized the results from “stage I” of that study (17,008 cases and 37,154 controls; 7,055,881 SNPs), except for those SNPs which were also analyzed in “stage II” (11,632 SNPs; using 8,572 AD cases and 11,312 controls in addition to those of stage I). For all analyses (JLIM, TWAS/FUSION, and the miRNA target gene set enrichment) we used IGAP stage I (i.e. genome-wide) summary statistics and for ADmiReQTL we used stage I data except for N=11,632 SNPs for which summary statistics from stage I and stage II combined were utilized instead. In addition, we obtained index SNPs from two more recent and larger AD GWAS (Jansen et al. (2018), Kunkle et al. (2018)); neither of these latter studies have made their summary statistics publicly available at the day of writing (June 2018). The first GWAS, by Jansen et al. (2018), utilized 24,087 cases and 55,058 controls and 9,862,738 SNPs in their discovery phase and 74,793 “AD by proxy” cases and 328,320 controls in their replication stage; altogether, they confirmed all but four of the IGAP loci and pinpointed an additional nine novel AD loci. In analyses using the ADmiReQTL workflow (see below), we used genome-wide significant index SNPs after meta-analysis of discovery and replication phase from their Supplementary Table 2. Kunkle et al. (2018) is a follow-up to the original IGAP report with sample increase of 29% for cases and of 13% for controls in stage I leading to overall 21,982 AD cases and 41,944 controls, and with overall 8,572 Alzheimer’s disease cases and 11,312 controls in stage 2 compared to the earlier report. In addition, they extended their stage I analyses to 9,456,058 SNPs while the number of variants in stage II remained the same as in the original paper. Overall, these updated analyses resulted in four new loci, one of which (i.e. *ECHDC3*) was also detected by Jansen et al. (2018). With ADmiReQTL, we used genome-wide significant index SNPs after meta-analysis of stage I and stage II data according to Table 1 of Kunkle et al. (2018).

### eQTL data

We performed a miRNA eQTL analysis using small RNA sequencing data of 345 samples of lymphoblastoid cell lines generated by the GEUVADIS consortium (Lappalainen et al. (2013)). In brief, this entailed genotype QC, expression quantification, batch correction, normalization and linear model fitting using Matrix eQTL (for details see Wohlers et al. (2018)). Overall, we combined the data from 468 miRNAs and 1,748,620 SNPs in our eQTL analyses resulting in a total of 3,191,687 association results. This resulted in 7,081 eQTLs at FDR of 0.05 (for 149 miRNAs and 5,658 SNPs). When defining p=0.01 for nominal significance, we observed 52,003 significant eQTLs (for 468 miRNAs and 45,911 SNPs).

### Data integration using ADmiReQTL

Statistical assessment of shared association signals between AD risk and miRNA expression using tools such as JLIM and TWAS/FUSION requires GWAS summary statistics, which are not yet available for the two latest and largest AD GWAS studies (Jansen et al. (2018), Kunkle et al. (2018)). However, it is still possible to assess novel loci identified by these studies by matching AD risk index SNPs to miRNA eQTLs within the implicated GWAS loci (defined as +/− 1 MB of the GWAS index SNP). To formalize this process, we developed a workflow (ADmiReQTL for “<u>AD</u>-<u>miR</u>NA-<u>eQTL</u> analysis pipeline”), which links each index GWAS SNP with eQTLs of all miRNAs within a specific locus (Figure 1). First, this entails assigning miRNA eQTL association data to the GWAS index SNP. For any given miRNA we then add the best, i.e. index, miRNA eQTL and its genomic information followed by computation of LD between GWAS SNP and eQTL index SNP; only variants with pairwise LD at r^2^≥0.1 are retained. Finally, and if available, the GWAS p-value of the best miRNA eQTL SNP is provided and compared to the best GWAS p-value in the region. As a result, ADmiReQTL comprehensively evaluates and enumerates all overlapping AD risk and miRNA expression association signals and filters those supported by LD.

**Figure 1:**
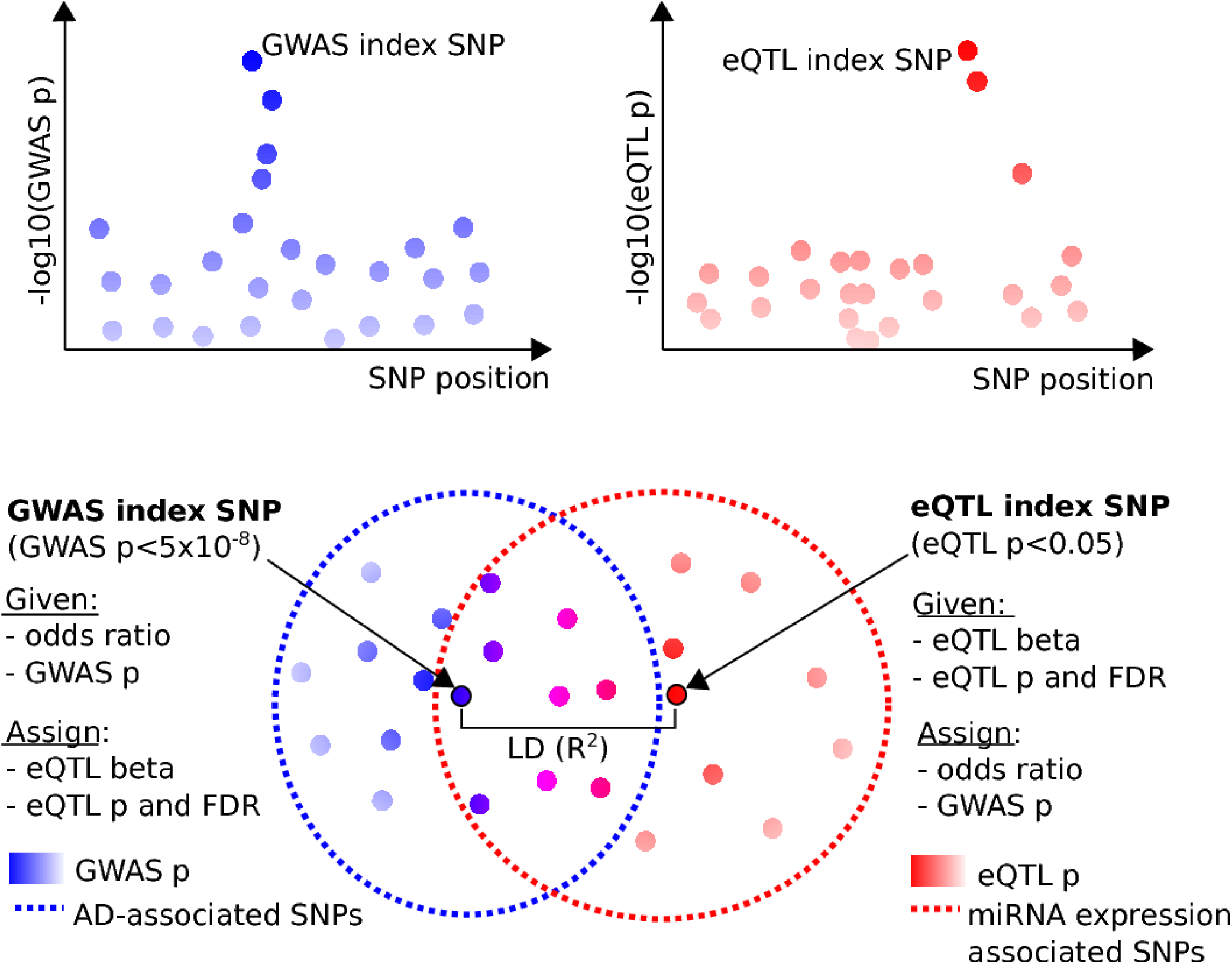
Illustration of the ADmiReQTL approach.

### miRNA target genes and miRNA target gene enrichment

Target genes were determined using two sources: First, from binding regions obtained from human brain AGO2 CLIP-Seq data published by Boudreau et al. (2014). Second, from miRTarBase, a database with a comprehensive listing of experimentally validated miRNA target genes (Chou et al. (2018). Enrichment of disease risk association was tested using PASCAL (Lamparter et al. (2016)) as described previously (Wohlers et al. (2018)).

### Application of JLIM software

JLIM is a recently developed statistical approach to assess whether signals in genetic association and eQTL data are linked by shared genetic effects. JLIM was run as described in the original publication (Chun et al. (2017), following instructions on the JLIM website (https://github.com/cotsapaslab/jlim). To ensure consistent genomic SNP coordinates among the eQTL and GWAS data, we lifted coordinates of the summary statistics SNPs to hg38 genome assembly using the PyLiftover library within Python.

### Application of TWAS/FUSION software

TWAS/FUSION is a recently developed tool to perform transcriptome-wide (here: miRNome-wide) association analyses by imputing miRNA expression into GWAS data from GWAS summary statistics and then testing whether expression is associated with disease risk. Here, we used TWAS/FUSION (Gusev et al. (2017)) as described on the TWAS/FUSION website (http://gusevlab.org/projects/fusion) using coordinates from the hg38 genome assembly. After consultation with the authors (i.e. Dr. Gusev; see acknowledgements), we updated required TWAS/FUSION reference files to hg38 and aligned SNP identifiers of the GWAS summary statistic with those used internally by TWAS/FUSION.

## 3 Results

In total, we assessed 65 index AD GWAS SNPs in 36 genomic regions, of which 28 SNPs in 15 regions were in proximity of an expressed miRNA and tested as eQTL for one or more miRNAs, see Supplementary Table 1. Our first analyses aimed at answering the question whether or not AD GWAS results (represented by summary statistics of the IGAP study) were linked to miRNA expression data (represented by the miRNA eQTL results that we generated from small RNA sequencing data in the GEUVADIS project) on a *genome-wide* level. To this end, we utilized JLIM, which assesses whether signals in genetic association and eQTL data are linked by shared genetic effects. However, these analyses revealed no significant loci with shared effects on a genome-wide scale (see below and Supplementary Table 2). Similarly, applying the TWAS/FUSION approach, which performs a miRNome-wide association study by predicting miRNA expression into GWAS data, did not reveal significant signals on a genome-wide level (see below and Supplementary Table 3). Taken together, these results do not suggest strong effects of AD genetic risk variants on miRNA expression in general, i.e. when analyses were based on the full genome-wide datasets with appropriate multiple testing correction. However, when individually matching the best miRNA eQTL SNPs within AD loci to AD GWAS index SNPs followed by filtering (ADmiReQTL, Supplementary Table 1), two miRNAs emerged, i.e. miR-6840-5p and miR-29c-5p. The expression of these two miRNAs may be regulated by variants showing genome-wide significant association with AD risk by GWAS (see Figures 2 and 3 and Subsection “Data integration using ADmiReQTL” for details).

**Figure 2:**
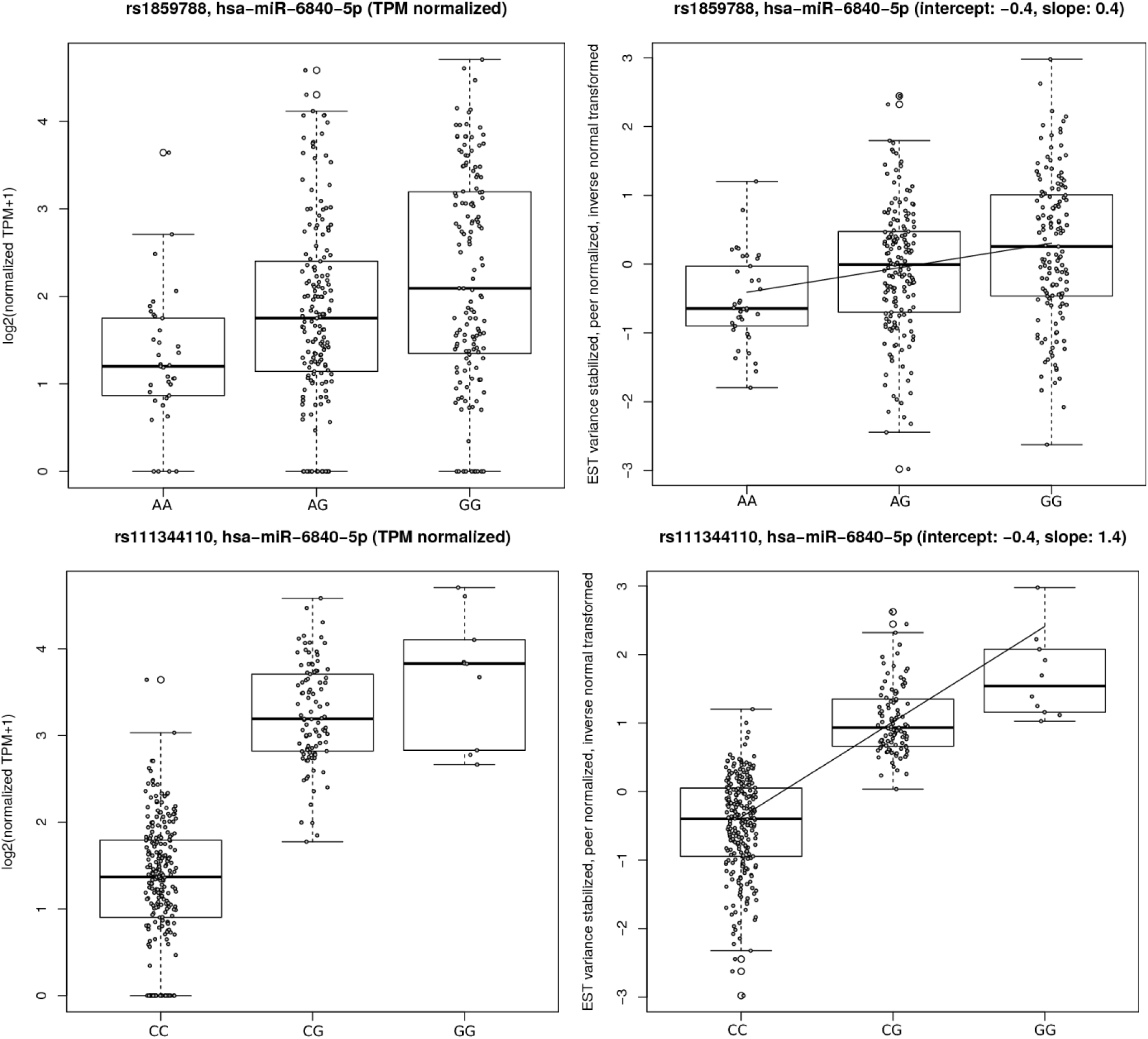
Boxplots of TPM normalized miRNA expression stratified by genotype (left) and of processed expression values used for eQTL computation (right). Top row: index variant. Bottom row: best eQTL SNP. Genotypes are ordered from hg38 reference allele (left) to alternative allele (right).

**Figure 3:**
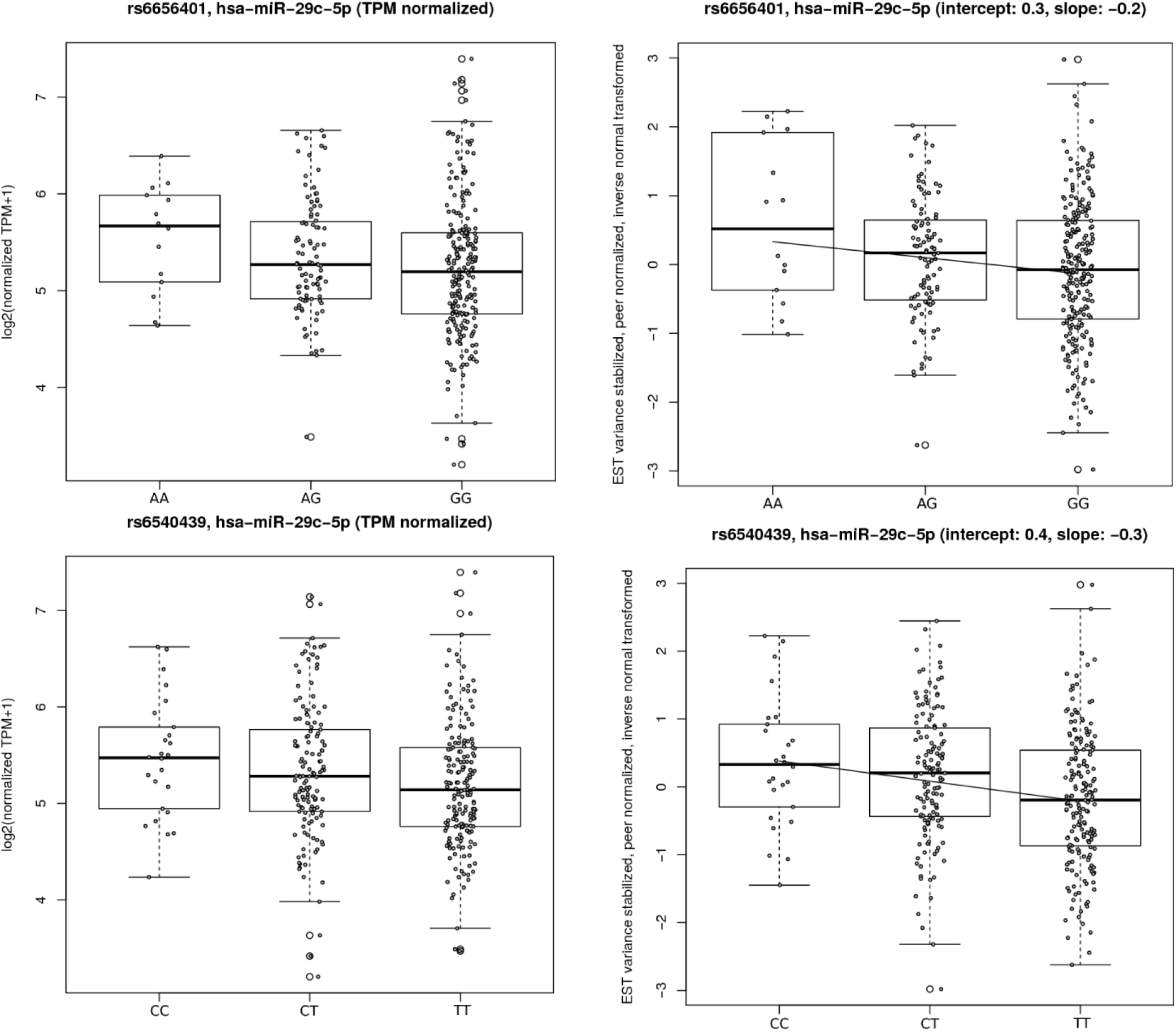
See caption of Figure 2.

### Application of JLIM and TWAS/FUSION

Neither JLIM nor TWAS/FUSION revealed any significant results when using FDR for multiple testing correction on the eQTL results indicating that there is no statistical support for a connection between the analyzed miRNA expression and AD GWAS data. For JLIM this overall negative result was indicated by the lack of nominal p-values less than 0.05 for any of the 13 loci and their overall 25 corresponding miRNAs (see Supplementary Table 2). Likewise, for TWAS/FUSION the lack of statistical support for AD association with miRNA expression was evidenced by the lack of FDR-corrected TWAS p-values less than 0.05 for any of the 49 miRNAs whose expression was sufficiently heritable for analysis (NB: miR-29c-5p could not be assessed; see Supplementary Table 3).

### Data integration using ADmiReQTL

The association with miR-6840-5p converges on the association of SNP rs1859788 with both AD risk (GWAS p=2.220E-15) and miRNA expression (eQTL p=1.582E-5; see Figure 2). This variant was reported as index AD GWAS SNP by Jansen et al. (2018) where it was assigned to the *NYAP1* locus (previously assigned to *ZCWPW1* by IGAP (2013)) on chromosome 7q22.1. The strongest miR-6840-5p eQTL SNP in the GEUVADIS sequencing data set is rs111344110 which is in only modest LD with the GWAS index SNP (r^2^ with rs1859788 = 0.108). This variant’s minor G-allele is correlated with the AD risk, major G-allele of rs1859788 and associated with increased miRNA expression (p-value of 5.279E-60; see Figure 2). The second highlighted miRNA is miR-29c-5p which is connected to AD genetics via SNP rs6656401 (GWAS p=5.692E-24, miRNA eQTL p=1.789E-2; see Figure 3). This variant was originally reported as genome-wide significantly associated with AD risk in the first IGAP GWAS and is located within the *CR1* locus at 1q32.2. This finding has since been replicated in many independent association studies, including the two most recent AD GWAS by Kunkle et al. and Jansen et al. SNP rs6656401 is located in an intron of the *CR1* gene but is also predicted to represent a gene regulatory variant according to Ensembl’s Variant Effect Predictor tool (VEP; McLaren et al. (2016); www.ensembl.org/Tools/VEP). The strongest miR-29c-5p eQTL in the GEUVADIS data is SNP rs6540439 (r^2^ with rs6656401 = 0.114) where it exhibits increased miRNA expression for the minor C allele, which is correlated with the AD risk, i.e. minor A-, allele of rs6656401 (p=5.295E-4; see Figure 3).

### miRNA target genes and miRNA target gene enrichment

Despite the observations from ADmiReQTL for miR6840-5p and miR-29c-5p, we observed no strong enrichment for significant GWAS signals in the target gene set of these two miRNAs using miRTarBase or human brain AGO2 CLIP-Seq data, with the exception of a weak nominal enrichment for miR-29c-5p in the latter data set (P=0.049) possibly driven by variants in *SORL1* (see Supplementary Table 4). We note that this lack of general enrichment of genetic association signals in the targets of either miRNA does not preclude functionally relevant effects in individual target genes. Among these there are at least two interesting candidates reported for miRNA-29c-5p, *BACE1* and *SORL1*.

## 4 Discussion

To the best of our knowledge, our study represents the first systematic assessment to investigate the potential overlap between genetics findings driving both risk for AD and miRNA expression. Despite using the largest hitherto available genome-wide datasets in each domain, we found no overarching strong evidence linking AD genetics to miRNA eQTLs. In other words, our findings provide a first indication that – perhaps with the exception of a few individual miRNAs – AD GWAS signals do not unfold their pathogenic effects by disturbing miRNA expression *in general*, at least in lymphoblast cell lines. This conclusion is different from that reached in similar studies with the aim to elucidate the impact of AD risk variants on mRNA expression. Several of these studies concluded that the predominant functional effect of many genetics variants underlying polygenic AD is to disturb mRNA expression (recently reviewed in Verheijen and Sleegers (2018)).

Despite utilizing large-scale “omics” data sets in our analyses, our study has some important limitations. First and foremost, the miRNA expression data utilized here was generated in human lymphoblastoid cell lines. MiRNA expression patterns in brain – especially in those regions (and cells) primarily affected by AD pathology – may be different from those found here. Second, and related to the previous point, only 468 miRNAs (out of 2588; Chou et al. (2018)) were found to be expressed at levels sufficient for analysis. It is therefore quite likely that our analyses missed to map potential eQTL effects simply owing to low abundance in lymphoblastoid cell lines while the same miRNAs may be expressed at higher levels in the brain. Neither of these limitations is addressable without access to sufficiently sized human brain miRNA expression data which currently do not exist. Third, with n=345 the data set used for the miRNA eQTL calculations is comparatively small and may have resulted in a loss in power for some aspects of our analyses. However, we note that the utilized data still originates from the largest currently available dataset with both genome-wide small-RNA and genotyping data generated in the same individuals.

Despite these limitations and the lack of an overall strong genetic effect on miRNA expression, we were able to pinpoint two “risk miRNA eQTLs” that may be of relevance in AD, i.e. GWAS SNPs correlating with levels of miR-29c-5p and miR-6840-5p. While none of the target gene sets of these miRNAs shows an enrichment of significant AD risk association signals in the IGAP GWAS dataset (see Results and Supplementary Table 4), it is interesting to note that previous data suggest that miR-29c can decrease the expression of beta-secretase 1 (BACE1; Lei et al., 2015), an enzyme involved in cleaving amyloid precursor protein (APP), which is of central relevance in AD pathogenesis (Selkoe and Hardy (2016)). Secondly, miR-29c-5p was also recently found to target *SORL1* mRNA in Ago2 CLIP-Seq experiments performed on human brain samples (Boudreau et al. (2014)). Genetically, *SORL1* has been linked to AD by both common (IGAP (2013)) and rare variant (Nicolas et al. (2016)) analysis, indicating that differential expression of this miRNA may exert some functionally relevant effects in AD pathogenesis. In conclusion, our large-scale integrative assessment provides only limited support for the hypothesis that AD GWAS variants act as miRNA eQTLs in lymphoblastoid cell lines on a large scale. Future work needs to confirm these findings in larger data sets, ideally assessing miRNA expression in human brain samples.

## Acknowledgements

Computational support was provided by the OMICS compute cluster at the University of Lübeck. We thank Dr. Chun for help with JLIM, Dr. Gusev for help with TWAS/FUSION, and Dr. Lill for advice on implementing the ADmiReQTL workflow.

## References

Boudreau RL, Jiang P, Gilmore BL, Spengler RM, Tirabassi R, Nelson JA, Ross CA, Xing Y, Davidson BL. Transcriptome-wide discovery of microRNA binding sites in human brain. Neuron. 2014;81(2):294–305.

Chou CH, Shrestha S, Yang CD, Chang NW, Lin YL, Liao KW, Huang WC, Sun TH, Tu SJ, Lee WH, Chiew MY, Tai CS, Wei TY, Tsai TR, Huang HT, Wang CY, Wu HY, Ho SY, Chen PR, Chuang CH, Hsieh PJ, Wu YS, Chen WL, Li MJ, Wu YC, Huang XY, Ng FL, Buddhakosai W, Huang PC, Lan KC, Huang CY, Weng SL, Cheng YN, Liang C, Hsu WL, Huang HD. miRTarBase update 2018: a resource for experimentally validated microRNA-target interactions. Nucleic Acids Res. 2018;46(D1):D296–D302.

Chun S, Casparino A, Patsopoulos NA, Croteau-Chonka DC, Raby BA, De Jager PL, Sunyaev SR, Cotsapas C. Limited statistical evidence for shared genetic effects of eQTLs and autoimmune-disease-associated loci in three major immune-cell types. Nat Genet. 2017;49(4):600–605.

Gusev A, Ko A, Shi H, Bhatia G, Chung W, Penninx BW, Jansen R, de Geus EJ, Boomsma DI, Wright FA, Sullivan PF, Nikkola E, Alvarez M, Civelek M, Lusis AJ, Lehtimäki T, Raitoharju E, Kähönen M, Seppälä I, Raitakari OT, Kuusisto J, Laakso M, Price AL, Pajukanta P, Pasaniuc B. Integrative approaches for large-scale transcriptome-wide association studies. Nat Genet. 2016;48(3):245–52.

International Genomics of Alzheimer’s Disease Project (IGAP). Meta-analysis of 74,046 individuals identifies 11 new susceptibility loci for Alzheimer’s disease. Nat Genet. 2013;45(12):1452–8.

Jansen I, Savage J, Watanabe K, Bryois J, Williams D, Steinberg S, Sealock J, Karlsson I, Hagg S, Athanasiu L, Voyle N, Proitsi P, Witoelar A, Stringer S, Aarsland D, Almdahl I, Andersen F, Bergh S, Bettella F, Bjornsson S, Braekhus A, Brathen G, de Leeuw C, Desikan R, Djurovic S, Dumitrescu L, Fladby T, Homan T, Jonsson P, Kiddle S, Rongve A, Saltvedt I, Sando S, Selbak G, Skene N, Snaedal J, Stordal E, Ulstein I, Wang Y, White L, Hjerling-Leffler J, Sullivan P, van der Flier W, Dobson R, Davis L, Stefansson H, Stefansson K, Pedersen N, Ripke S, Andreassen O, Posthuma D. Genetic meta-analysis identifies 9 novel loci and functional pathways for Alzheimers disease risk. bioRxiv 258533; doi: https://doi.org/10.1101/258533.

Kunkle BW, Grenier-Boley B, Sims R, Bis JC, Naj AC, Boland A, Vronskaya M, Van der Lee SJ, Amlie-Wolf A, Bellenguez C, Frizatti A, Chouraki V, Alzheimer’s Disease Genetics Consortium (ADGC), European Alzheimer’s Disease Initiative (EADI), Cohorts for Heart and Aging Research in Genomic Epidemiology Consortium (CHARGE), Genetic and Environmental Risk in Alzheimer’s Disease Consortium (GERAD/PERADES), Schmidt H, Hakonarson H, Munger R, Schmidt R, Farrer LA, Van Broeckhoven C, O’Donovan MC, Destefano AL, Jones L, Haines JL, Deleuze J, Owen MJ, Gudnason V, Mayeux RP, Escott-Price V, Psaty BM, Ruiz A, Ramirez A, Wang L, van Duijn CM, Holmans PA, Seshadri S, Williams J, Amouyel P, Schellenberg GD, Lambert J, Pericak-Vance MA. Meta-analysis of genetic association with diagnosed Alzheimer’s disease identifies novel risk loci and implicates Abeta, Tau, immunity and lipid processing. bioRxiv 294629; doi: https://doi.org/10.1101/294629.

Lamparter D, Marbach D, Rueedi R, Kutalik Z, Bergmann S. Fast and Rigorous Computation of Gene and Pathway Scores from SNP-Based Summary Statistics. PLoS Comput Biol. 2016;12(1):e1004714.

Lappalainen T, Sammeth M, Friedländer MR, ‘t Hoen PA, Monlong J, Rivas MA, Gonzàlez-Porta M, Kurbatova N, Griebel T, Ferreira PG, Barann M, Wieland T, Greger L, van Iterson M, Almlöf J, Ribeca P, Pulyakhina I, Esser D, Giger T, Tikhonov A, Sultan M, Bertier G, MacArthur DG, Lek M, Lizano E, Buermans HP, Padioleau I, Schwarzmayr T, Karlberg O, Ongen H, Kilpinen H, Beltran S, Gut M, Kahlem K, Amstislavskiy V, Stegle O, Pirinen M, Montgomery SB, Donnelly P, McCarthy MI, Flicek P, Strom TM; Geuvadis Consortium, Lehrach H, Schreiber S, Sudbrak R, Carracedo A, Antonarakis SE, Häsler R, Syvänen AC, van Ommen GJ, Brazma A, Meitinger T, Rosenstiel P, Guigó R, Gut IG, Estivill X, Dermitzakis ET. Transcriptome and genome sequencing uncovers functional variation in humans. Nature. 2013;501(7468):506–11.

Lei X, Lei L, Zhang Z, Zhang Z, Cheng Y. Downregulated miR-29c correlates with increased BACE1 expression in sporadic Alzheimer’s disease. Int J Clin Exp Pathol. 2015;8(2):1565–74.

McLaren W, Gil L, Hunt SE, Riat HS, Ritchie GR, Thormann A, Flicek P, Cunningham F. The Ensembl Variant Effect Predictor. Genome Biology 2017(1):122.

Nicolas G, Charbonnier C, Wallon D, Quenez O, Bellenguez C, Grenier-Boley B, Rousseau S, Richard AC, Rovelet-Lecrux A, Le Guennec K, Bacq D, Garnier JG, Olaso R, Boland A, Meyer V, Deleuze JF, Amouyel P, Munter HM, Bourque G, Lathrop M, Frebourg T, Redon R, Letenneur L, Dartigues JF, Génin E, Lambert JC, Hannequin D, Campion D; CNR-MAJ collaborators. SORL1 rare variants: a major risk factor for familial early-onset Alzheimer’s disease. Mol Psychiatry. 2016;21(6):831–6.

Selkoe DJ, Hardy J. The amyloid hypothesis of Alzheimer’s disease at 25 years. EMBO Mol Med. 2016;8(6):595–608.

Verheijen J and Sleegers K. Understanding Alzheimer Disease at the Interface between Genetics and Transcriptomics. Trends Genet. 2018;34(6):434–447.

Wohlers I, Bertram L, Lill CM. Evidence for a potential role of miR-1908-5p and miR-3614-5p in autoimmune disease risk using genome-wide analyses. bioRxiv 286260; doi: https://doi.org/10.1101/286260.

